# Estimating the effect of a scanner upgrade on measures of grey matter structure for longitudinal designs

**DOI:** 10.1101/2020.08.28.271296

**Authors:** Evelyn Medawar, Ronja Thieleking, Iryna Manuilova, Maria Paerisch, Arno Villringer, A. Veronica Witte, Frauke Beyer

## Abstract

Longitudinal imaging studies are crucial for advancing the understanding of brain development over the lifespan. Thus, more and more studies acquire imaging data at multiple time points or with long follow-up intervals. In these studies changes to magnetic resonance imaging (MRI) scanners often become inevitable which may decrease the reliability of the MRI assessments and introduce biases.

We therefore investigated the difference between MRI scanners with subsequent versions (3 Tesla Siemens Verio vs. Skyra fit) on the cortical and subcortical measures of grey matter in 116 healthy, young adults using the well-established longitudinal FreeSurfer stream for T1-weighted brain images. We found excellent between-scanner reliability for cortical and subcortical measures of grey matter structure (intra-class correlation coefficient > 0.8). Yet, paired t-tests revealed statistically significant differences in at least 75% of the regions, with percent differences up to 5%, depending on the outcome measure. Offline correction for gradient distortions only slightly reduced these biases. Further, T1-imaging based quality measures systematically differed between scanners.

We conclude that scanner upgrades during a longitudinal study introduce bias in measures of cortical and subcortical grey matter structure. Therefore, before upgrading a MRI scanner during an ongoing study, researchers should prepare to implement an appropriate correction method for these effects.

## 2 Introduction

Many longitudinal neuroimaging studies of aging and development investigate changes in local grey matter volume (GMV) over time to identify biomarkers relevant to health and disease. Notably, in the past decade many large-scale studies have implemented longitudinal designs in the general population (with at least two timepoints: Bycroft et al. (2018); Ikram et al. (2015), second timepoint currently being acquired: Loeffler et al. (2015); Bamberg et al. (2015)).

Such longitudinal imaging studies assess within-subject differences and thereby benefit from reduction of error variance and confounding. Yet, scanner changes often become inevitable with long follow-up intervals (4-6 years) in these studies, entailing issues of reliability because of changes in signal-to-noise ratio or image intensity (Preboske et al. 2006; Takao et al. 2010; Ewers et al. 2006; Chen et al. 2014). This is especially problematic in the case of two-visit longitudinal imaging studies where measurement occasion may be collinear with scanner upgrade, making it difficult to draw unbiased conclusions on within-subject change. In contrast, scanner upgrades will affect cross-sectional designs less as scanner version can be modelled like a site effect (Fortin et al. 2018).

Before the follow-up of the LIFE-Adult Study, a two-visit longitudinal imaging study with a long inter-visit interval (5-7 years), we had to decide on the upgrade of the study scanner from Siemens Verio to Skyra fit (Loeffler et al. 2015). At the time (end of 2017), most studies on the effects of scanner upgrades had investigated small samples (n<15) or voxel-based morphometry estimates of grey matter (GM) structure, with varying estimates of reliability and bias (Jovicich et al. 2009; Shuter et al. 2008; Takao, Hayashi, and Ohtomo 2013). Thus, the impact of a scanner upgrade on region- and vertex-wise measures of cortical GM (thickness, area and volume) as well as subcortical GM volume still lacked quantification. Also, these studies did not take into account gradient distortion correction which has been shown to partly account for variation between scanners (Jovicich et al. 2006; Cannon et al. 2014).

Here, we therefore investigated the difference between scanners with subsequent versions (3 Tesla Siemens Verio vs. Skyra fit) on the cortical and subcortical measures of GM in a large sample of healthy, young adults. Differences between the systems included the changes introduced by software and hardware upgrades (update to syngo MR E11 software, Tim 4G body coil, installation of DirectRF) and systematic differences due to B0 and B1 fields. Because we were about to decide on the upgrade (which we eventually declined), we could not perform a comparison of the same scanner pre/post upgrade.

Using the validated longitudinal FreeSurfer stream, we expected the reliability of whole-brain and regional GM measures to be similar to previous studies investigating between-site reliability (Reuter et al. 2012; Keshavan et al. 2016; Jovicich et al. 2013). Based on previous upgrade studies, we hypothesized a systematic bias with varying effect sizes and direction in cortical and subcortical regions (Jovicich et al. 2009; Han et al. 2006). Finally, we expected gradient distortion correction to improve reliability and reduce bias.

## 3 Methods

### 3.1 Sample

121 healthy participants (age in years: mean = 30.02, sd = 8.24; 61 females) were scanned on two different 3 Tesla MRI scanners with subsequent versions (Magnetom Verio syngo MR B17, Magnetom Skyra fit syngo MR E11 (Siemens, Erlangen)). Due to a pending version update of the Verio scanner, all participants were first scanned at the Verio and then at the Skyra scanner. On average, 2.14 months (sd = 1.09 months) passed in-between sessions.

5 participants did only participate in the first scanning session at the Verio and were therefore excluded in the following analysis. The study was approved by the local ethics committee at the University of Leipzig and all participants gave written informed consent according to the Declaration of Helsinki.

### 3.2 Imaging sequence

On both scanners, anatomical T1-weighted imaging was performed with a magnetization-prepared rapid gradient-echo (MPRAGE) sequence (TR=2300 ms,TE=2.98 ms, TI=900 ms, acceleration: GRAPPA factor 2, flip angle: 9°, imaging matrix 256 x 240 x 176 and voxel size= 1 *mm*^3^, with prescan normalize option) according to the ADNI protocol (Jack et al. 2011). Additional sequences, i.e. diffusion-weighted imaging, were also acquired and are discussed elsewhere (Thieleking et al., in prep). On the Skyra scanner, both online 3D gradient distortion-corrected images (“D”) and images not corrected for distortions (“ND”) were available. The Verio scanner delivered the images without gradient-distortion correction (“ND”).

In the main analysis, we compared the Verio “ND” and the distortion-corrected “D” Skyra data, in the gradient distortion correction analysis we compared the offline `gradunwarp` distortion corrected Skyra ND and Verio ND (see the diagram of the study flow in Figure 1).

**Figure 1:**
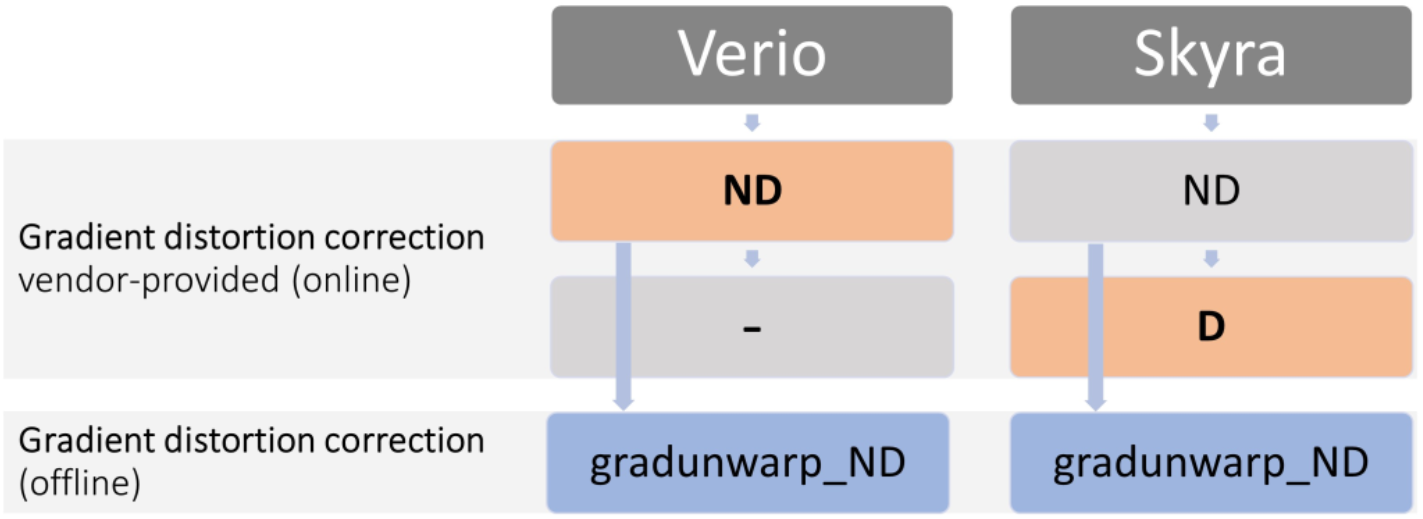
Overview of the scanner outcome images and the performed analyses. Orange: input images for main analysis based on standard output provided by scanner (Verio without correction, Skyra with distortion correction). Blue: input images for secondary analysis based on original images from scanner and subsequent offline distortion correction with gradunwarp.

### 3.3 Preprocessing

#### 3.3.1 Gradient distortion correction

Gradient distortion correction has been shown to contribute to measurement error in repeated sessions of anatomical brain imaging (Takao et al. 2010). Accordingly, correcting for distortion correction can improve the reproducibility of intensity data significantly (Jovicich et al. 2006). For the Verio scanner, the vendor provided no online distortion correction while the Skyra system offered online 3D-distortion correction. To assess the effect of this processing step on reliability and bias, we applied an identical tool for offline gradient distortion correction on the ND sequences from both scanners.

Gradient unwarping calculates the geometric displacement based on the spherical expansion of the magnetic gradient fields and applies it to the image. We used the gradunwarp implementation [(https://github.com/Washington-University/gradunwarp)] in Python 2.7. We visually compared the original and the `gradunwarp` result files to determine the appropriate number of sampling points and interpolation order. Based on this, we chose 200 sampling points and 4th order interpolation (`--fovmin −0.2 --fovmax 0.2 --numpoints 200 --interp_order 4`) because this yielded most similar intensity distributions. After unwarping, we repeated FreeSurfer’s cross-sectional and longitudinal stream for these images. Then, we assessed the reliability and bias in cortical and subcortical ROI measures between the `gradunwarp` distortion corrected Skyra ND and Verio ND images.

#### 3.3.2 FreeSurfer analysis

To extract reliable volume and thickness estimates, we processed the T1-weighted images with the longitudinal stream in FreeSurfer (Reuter et al. 2012). Within this pipeline, an unbiased within-subject template space is created using robust, inverse consistent registration (Reuter, Rosas, and Fischl 2010; Reuter and Fischl 2011). The longitudinal stream increases the reliability of cortical and subcortical GM estimates compared to the cross-sectional stream and is thus appropriate for longitudinal studies (Jovicich et al. 2013). We used FreeSurfer version 6.0.0p1 with the default parameters `recon-all -all -parallel -no-isrunning -openmp 8`, which include non-parametric non-uniform intensity normalization with the MINC tool nu_correct. First, we ran the recon-all longitudinal stream for Verio ND and Skyra D. Then, we repeated this analysis for the gradient-unwarped Verio and Skyra ND T1-weighted images.

#### 3.3.3 Quality Assessment

We visually checked the cross-sectional as well as the longitudinal runs for errors in white matter segmentation and misplaced pials (Klapwijk et al. 2019). There were 17 cases where the pial surface expanded into nonbrain tissue. These were corrected by either editing the brainmask in the longitudinal template or by correcting the cross-sectional runs. After correction, we re-ran the longitudinal template creation step and the longitudinal timepoints. No issues regarding white matter segmentation were noticed.

To quantify potential differences in image quality between scanners, we compared different quality control measures provided by `mriqc` version 0.15.0 (Esteban et al. 2017). We used coefficient of joint variation (CJV) which was highlighted as an important predictor of image quality in (Esteban et al. 2017). Furthermore, we compared contrast-to-noise ratio (CNR) to quantify the difference between grey and white matter intensity distributions and the entropy focus criterion (EFC) to describe the amount of ghosting and blurring induced by head motion. We performed `mriqc` on the Verio ND, Skyra ND and Skyra D images.

#### 3.3.4 Outcomes

As outcomes we selected cortical thickness (CT), area (CA) and volume (CV) estimates for regions of interests defined by the Desikan-Killiany (DK) cortical parcellation (64 ROIs for both hemispheres). Subcortical volumes were extracted from FreeSurfer’s subcortical segmentation (“aseg.mgz”, 18 bilateral ROIs). We analyzed all ROIs per hemisphere. Subcortical volumes were not adjusted for head size because during the longitudinal stream, both images are normalized to the same head size.

### 3.4 Analysis

All statistical analysis were performed in R version 3.6.1 (R Core Team 2017).

#### 3.4.1 Reliability and percent difference of cortical and subcortical GM measures

To assess the reliability of the GM estimates, we calculated the intra-class correlation coefficient (ICC) using the two-way mixed effect ICC model for single measures with absolute agreement (Shrout and Fleiss 1979), implemented in the package psy. The ICC yields a value between 0 and 1, with a similar interpretation as the Pearson’s correlation coefficient, but takes into account a possible bias between rater (i.e. scanners). We calculated ICC for each cortical DK and subcortical ROI and reported the estimate and 95% confidence interval, derived by bootstrapping. According to (Cicchetti 1994), we considered an ICC below .4 to be poor, between .40 and .59 to be fair; .60 and .74 to be good and between .75 and 1.00 to be excellent. In order to assess the relative difference of GM measures between scanners, we calculated percent difference (PD) (also termed variability error (Jovicich et al. 2013; Iscan et al. 2015)). We calculated the mean of the PD for each ROI *j* across *n* participants according to

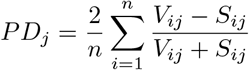

where *V_ij_* is the GM measure of a ROI measured on the Verio, *S_ij_* is the GM measure of a ROI measured on the Skyra.

Finally, we performed paired t-tests to inform about the direction and statistical significance of potential systematic differences between scanners. Here, we used Benjamini-Hochberg correction to adjust p-values per cortical GM measure and deemed differences to be significant at *p_adj_* <0.05 (Benjamini and Hochberg 1995). We reported T-value, uncorrected and corrected p-values.

We compared the improvement induced by using the same unwarping procedure for both Skyra ND and Verio ND images by applying paired t-tests to the ICC and PD measures of CT and subcortical volume from the secondary analysis (`gradunwarp` skyra D, Verio D) and the original analysis (Skyra D, Verio ND).

#### 3.4.2 Vertex-wise estimation of reliability and percent difference

For whole-brain visualization, we performed vertex-wise calculations on the fsaverage template following (Liem et al. 2015) in Matlab version 9.7 (2019b). We calculated ICC and PD for cortical thickness, area and volume to visualize reliability and difference between scanners on a vertex-wise level.

#### 3.4.3 Quality metrics

For the quality metrics from `mriqc`, we used linear mixed models (LMM) to assess differences between scanners (Verio, Skyra) and acquisitions (D, ND) using `lmerTest`. Significance was defined based on model comparisons (using Chi-square test with R’s anova) between LMM including either scanner or acquisition as a fixed effect and null models only including the random effects of subject. Significance was defined as p < 0.05. We also tested whether CNR was associated with regional CT, independent of scanner, using a LMM with both factors. We reported *β* estimates, raw and Benjamini-Hochberg adjusted p-values.

#### 3.4.4 Data and Code availability

Region of interest data and code used for this publication are available on github (https://github.com/fBeyer89/life_upgrade). Under certain conditions, the authors may also provide access to the MRI data.

## 4 Results

### 4.1 Differences in cortical GM measures between scanners

Figures 2, 3 and 4 summarize the results for CT, CA and CV, respectively.

**Figure 2:**
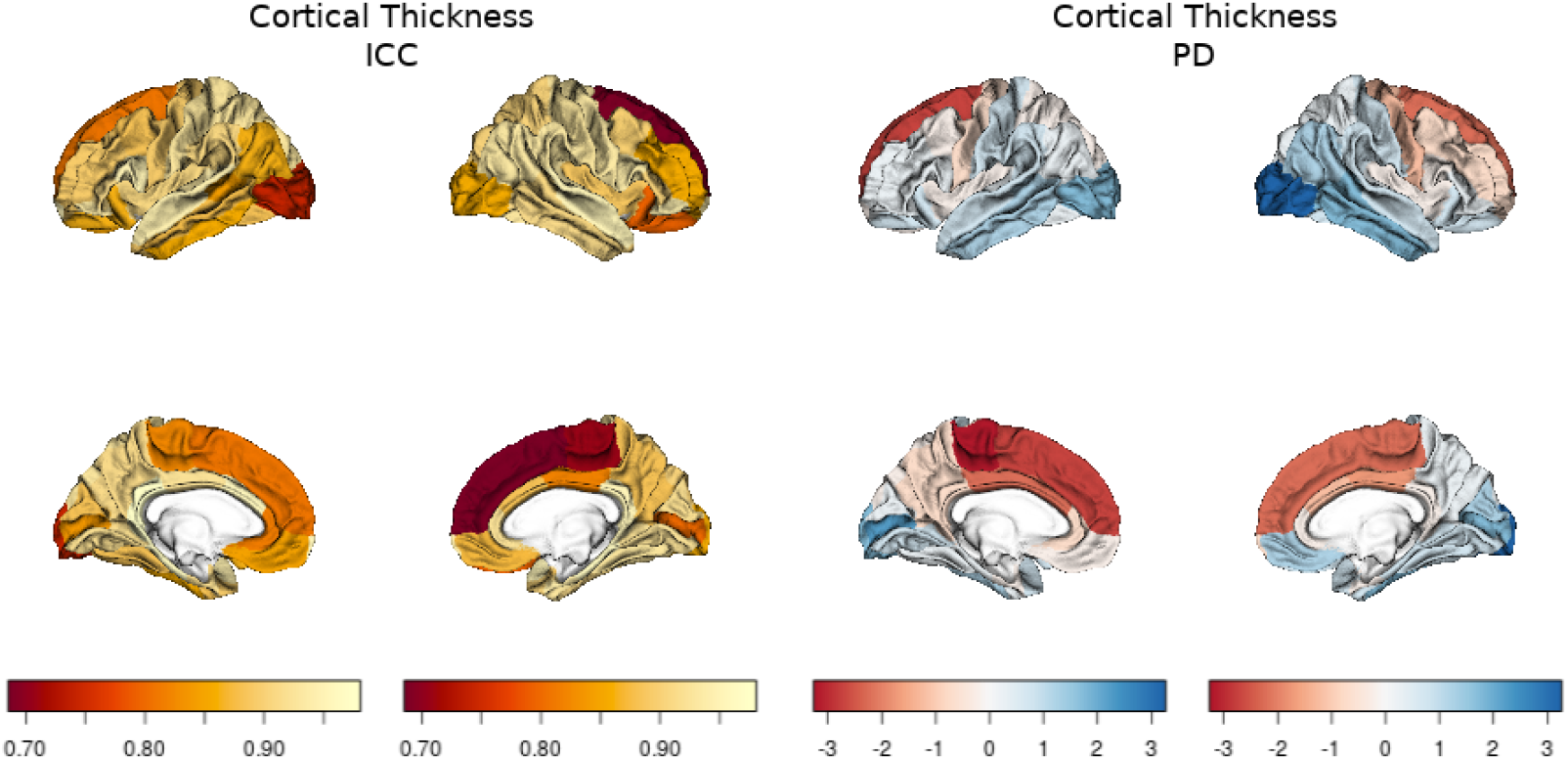
left panel: CT ICC, right panel: CT PD (for each panel, left column shows lateral and medial view of left hemisphere, right column shows lateral and medial view of right hemisphere), negative values:Skyra>Verio, positive values: Verio>Skyra

**Figure 3:**
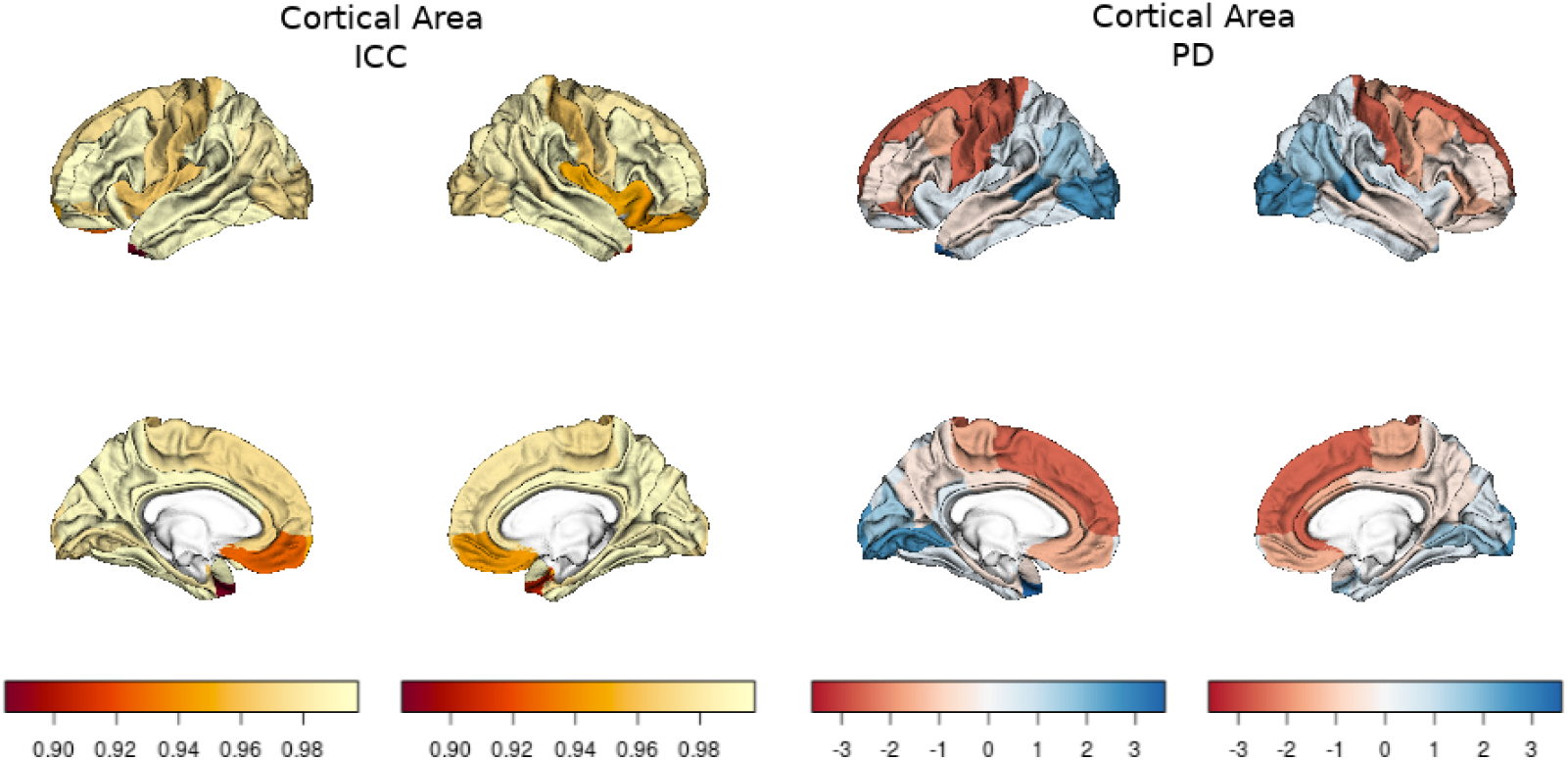
left panel: CA ICC, right panel: CA PD (for each panel, left column shows lateral and medial view of left hemisphere, right column shows lateral and medial view of right hemisphere), negative values:Skyra>Verio, positive values: Verio>Skyra

**Figure 4:**
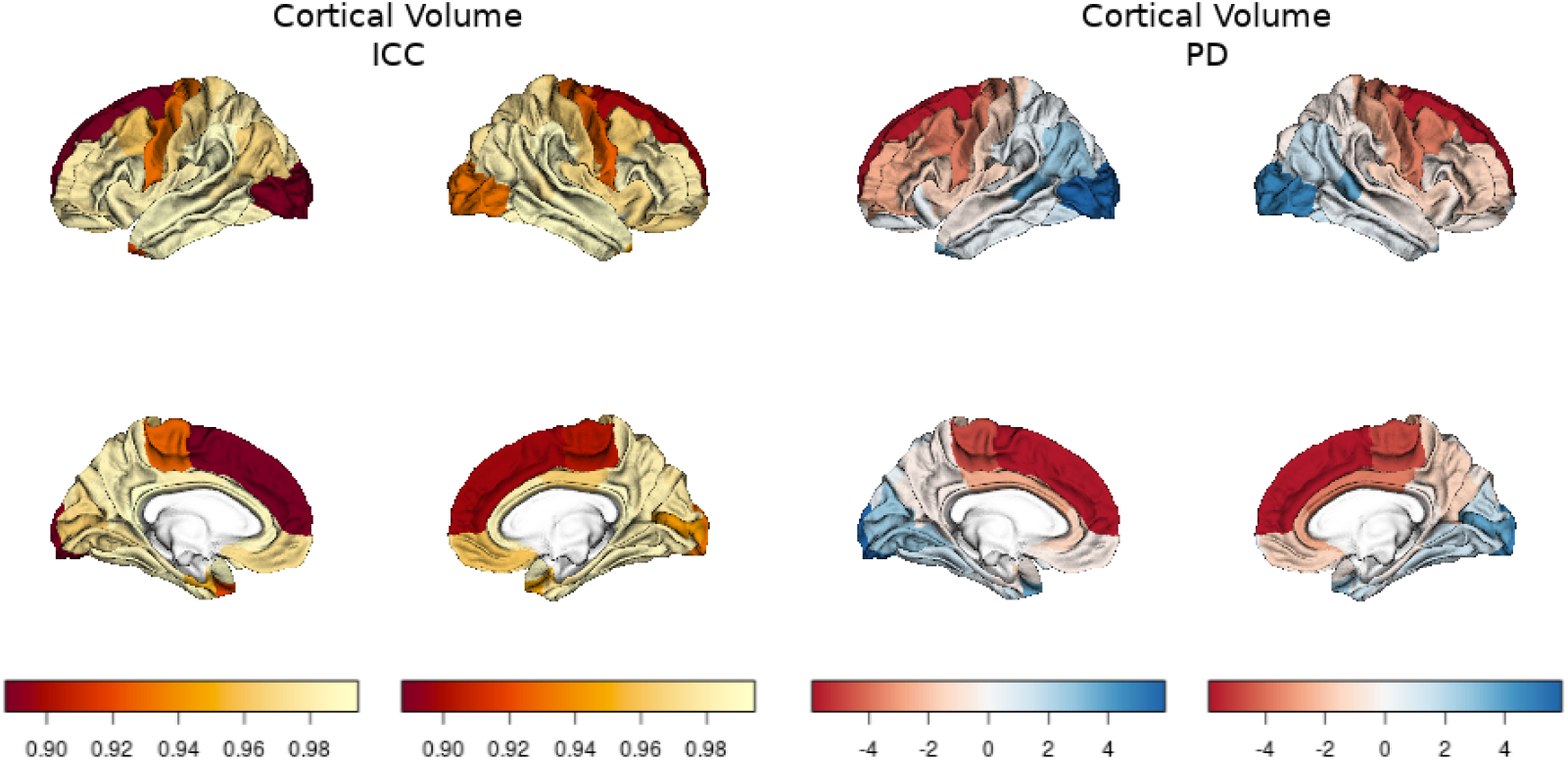
left panel: CV ICC, right panel: CV PD (for each panel, left column shows lateral and medial view of left hemisphere, right column shows lateral and medial view of right hemisphere), negative values:Skyra>Verio, positive values: Verio>Skyra

Overall, the ICC or scan-rescan reliability was excellent (CT: mean=0.89, min=0.69, max=0.98; CA: mean=0.98, min=0.88, max=1; CV: mean=0.97, min=0.89, max=0.99). The PD was around 2-4% for CT and CA (CT: mean=0.13, min=-3.16, max=3.23; CA: mean=-0.17, min= −2.58, max=3.56), and slightly higher for CV (CV: mean=-0.43, min=-5.38, max=5.84). Most pronounced differences were located in medial and lateral frontal and central regions, where CT, CA and CV were lower in Verio compared to Skyra. Higher values for CT, CA and CV for Verio compared to Skyra were found in lateral occipital, inferior and middle temporal regions. Overall, the bias direction seems to follow a frontal-to-occipital pattern. Accordingly, paired t-tests indicated systematic differences between scanners for most regions of interest (FDR-corrected, CT: 75% of all 64 bilateral ROIs, CA: 92.2%, CV: 81.2%). For detailed results per cortical region see Tables 1.1, 2.1 and 3.1 in the Supplementary Material. The vertex-wise analysis showed similar effects of a frontal-to-occipital pattern, with higher CT, CA and CV in the central gyrus, and in the gyri of the temporal lobe (regions: postcentral, superiortemporal, inferiortemporal) for Verio compared to Skyra (also see Supplementary Material, Figures 1.2 - 3.3).

### 4.2 Differences in subcortical measures between scanners

As shown in Table 1, subcortial regions, similar to cortical areas, showed excellent reliability for all ROI (mean=0.95, min=0.81, max=0.99). The PD was around 2-4% (mean=2.77%, min=1.22%, max=9.52%), with an exceptionally high value of 9.5% for left Accumbens. Significant differences between scanners were evident for most regions (FDR-corrected, 85.7% of all 14 bilateral ROIs).

**Table 1:** Reliability and percent differences for subcortical volumes (T<0 reflects Skyra>Verio, T>0 reflects Verio>Skyra)

| ROI | hemi | ICC | lower ICC | upper ICC | PD | T | p | adj.p |
| --- | --- | --- | --- | --- | --- | --- | --- | --- |
| Thalamus | Left | 0.97 | 0.96 | 0.98 | 1.90 | -12.27 | 0.00 | **0** |
| Thalamus | Right | 0.98 | 0.97 | 0.98 | 1.61 | -9.37 | 0.00 | **0** |
| Caudate | Left | 0.99 | 0.99 | 0.99 | 1.65 | -10.64 | 0.00 | **0** |
| Caudate | Right | 0.98 | 0.98 | 0.99 | 2.12 | -15.72 | 0.00 | **0** |
| Putamen | Left | 0.98 | 0.98 | 0.99 | 1.58 | -6.22 | 0.00 | **0** |
| Putamen | Right | 0.99 | 0.99 | 0.99 | 1.22 | -4.85 | 0.00 | **0** |
| Pallidum | Left | 0.96 | 0.95 | 0.97 | 2.48 | -0.44 | 0.66 | 0.66 |
| Pallidum | Right | 0.95 | 0.93 | 0.96 | 2.53 | -3.58 | 0.00 | **0** |
| Hippocampus | Left | 0.96 | 0.95 | 0.96 | 2.01 | -8.51 | 0.00 | **0** |
| Hippocampus | Right | 0.97 | 0.97 | 0.98 | 1.61 | -8.16 | 0.00 | **0** |
| Amygdala | Left | 0.91 | 0.89 | 0.93 | 3.57 | -1.47 | 0.14 | 0.15 |
| Amygdala | Right | 0.94 | 0.94 | 0.95 | 3.00 | -2.73 | 0.01 | **0.01** |
| Accumbens | Left | 0.81 | 0.70 | 0.85 | 9.52 | -9.30 | 0.00 | **0** |
| Accumbens | Right | 0.95 | 0.93 | 0.96 | 3.91 | -5.53 | 0.00 | **0** |

### 4.3 QA measures

First, we compared CJV, CNR and EFC, three quality measures from mriqc between Verio ND and Skyra D acquisitions. We aimed to determine whether differences in basic signal properties might underlie the observed differences in measures of GM structure.

We found that overall Verio ND T1-weighted images had higher CNR (*β*=0.15, p < 0.001), lower EFC (*β*= −0.04, p < 0.001) and lower CJV (*β*=-0.02, p < 0.001) compared to the Skyra D images, also see Figure 5. This indicates overall better data quality on the Verio scanner.

**Figure 5:**
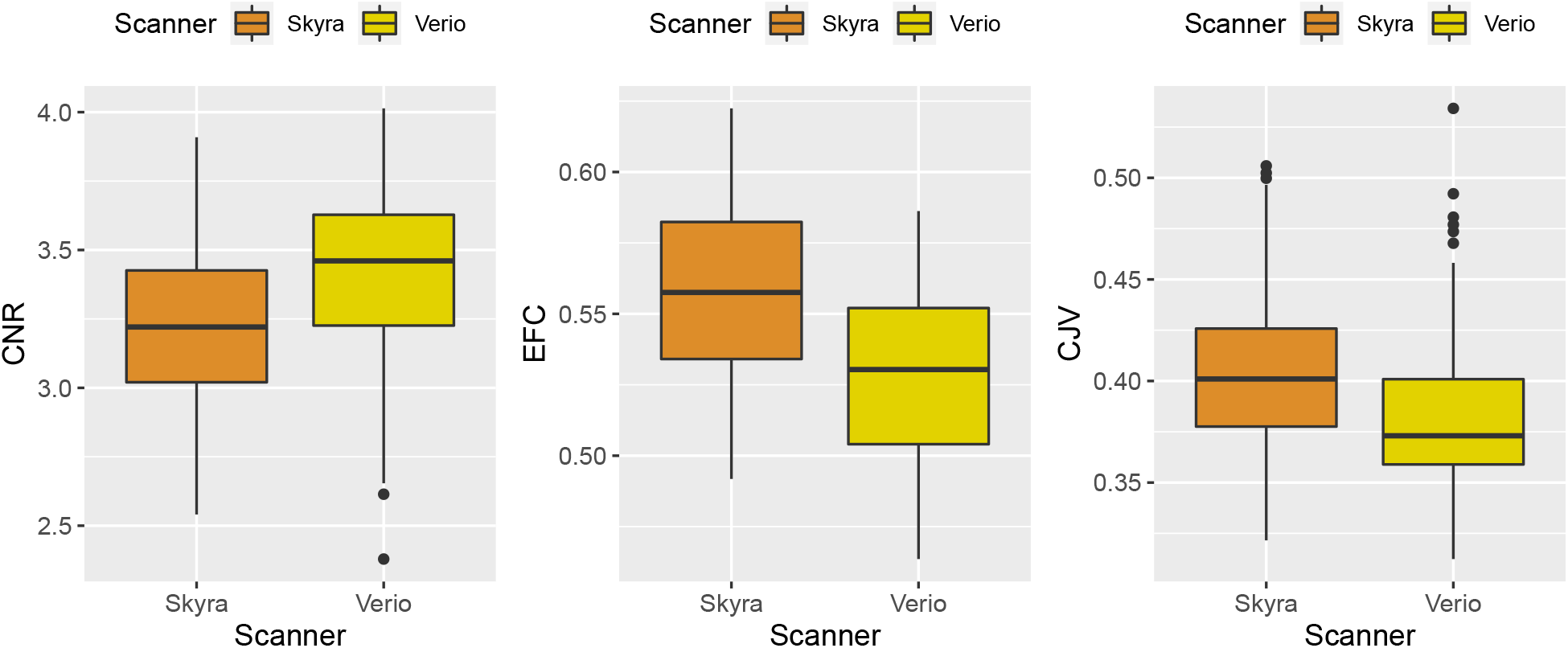
Quality metrics (CNR (left panel), EFC (middle panel) and CJV (right panel)) compared between Skyra D (orange) and Verio ND (yellow) acquisitions, showing overall higher data quality on the Verio scanner

When comparing the acquisitions with and without vendor-provided online gradient distortion correction on the Skyra scanner (“D” and “ND”), we observed that the distortion correction increased CNR (*β*=-0.113, p < 0.001, see Figure 6, left panel). When only considering ND acquisitions from both scanners, we also see higher CNR on the Verio scanner (*β*=0.088, p < 0.001, see Figure 6, right panel). This indicates that other factors than distortion correction underlie higher CNR on the Verio scanner.

**Figure 6:**
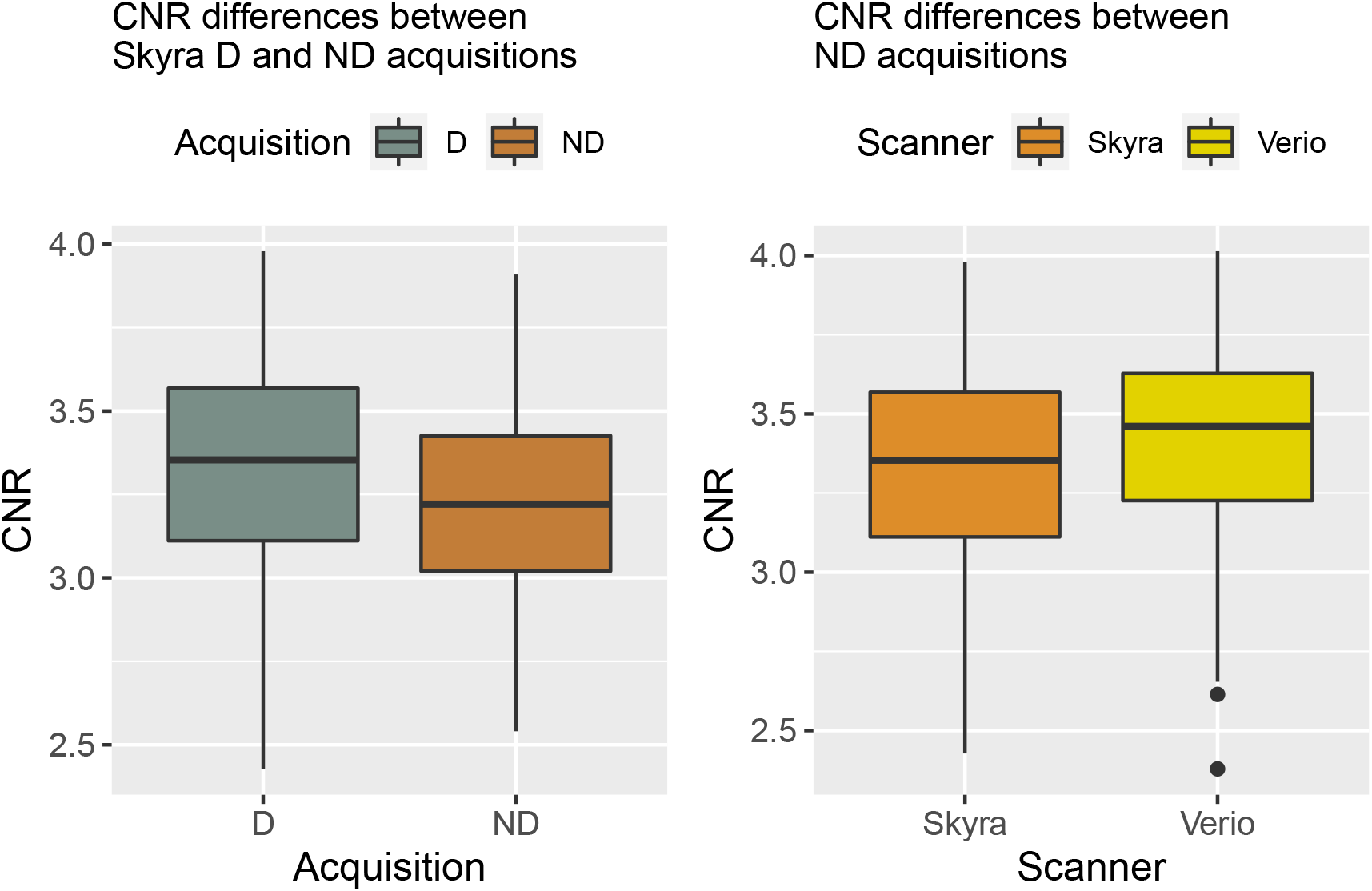
CNR differences between the Skyra ND and D acquisitions (left panel) and between Skyra and Verio ND acquisitions (right panel), showing higher CNR irrespective of gradient distortion on the Verio scanner

Similar to (Shuter et al. 2008), we investigated whether increased CNR would predict differences in CT. Here, we found that higher CNR across both scanners was associated with higher CT for most regions (see Figure 7, left panel). Moreover, scanner predicted CT independent of CNR in the same regions as shown above (see Figure 7, right panel, and Table 4.1 in the Supplementary Material).

**Figure 7:**
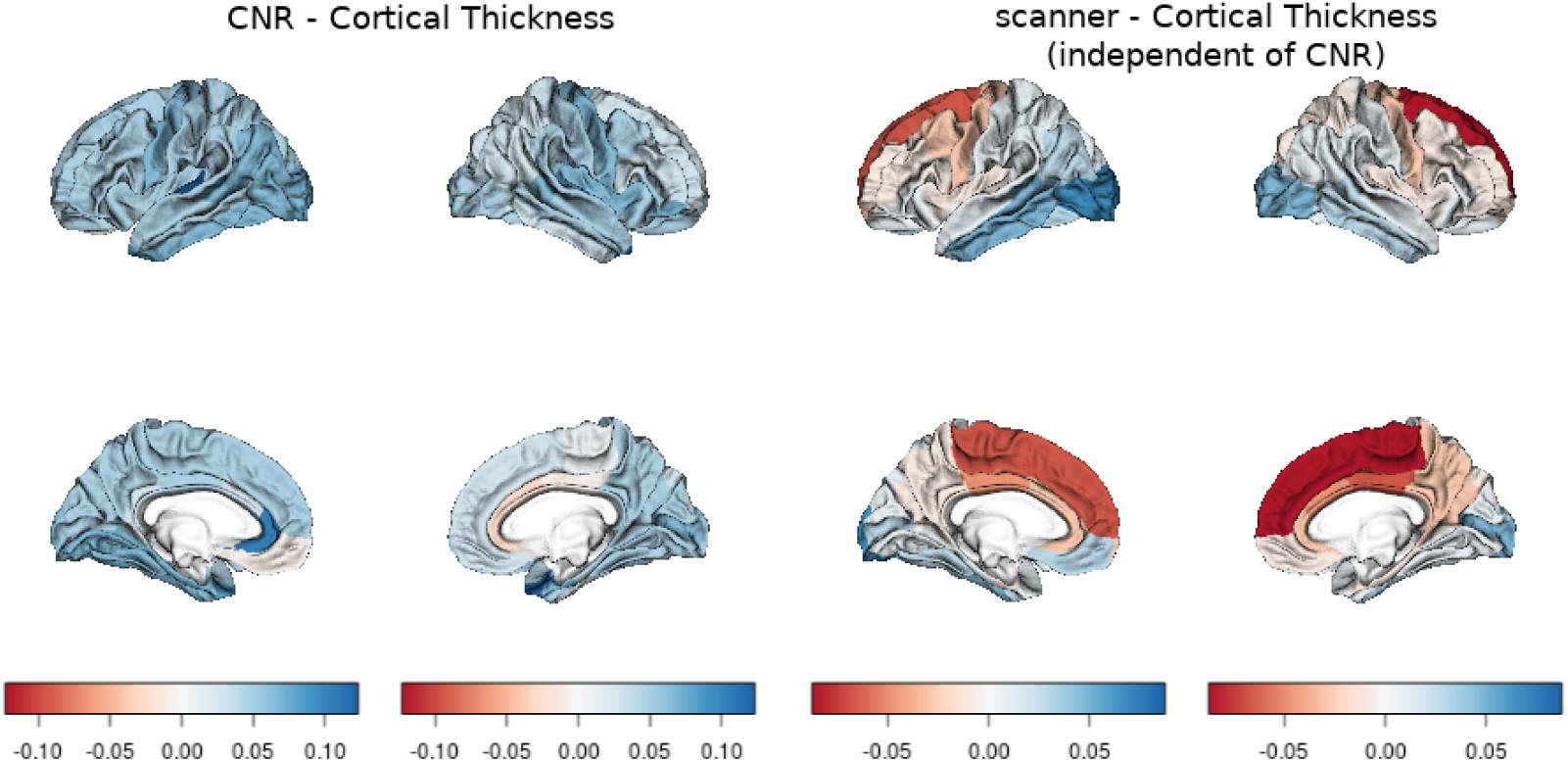
Association of CNR (left panel) and scanner (right panel, negative values indicate Skyra>Verio) with cortical thickness, shown as coefficients from a linear mixed model including both terms. Left column shows lateral and medial view of left hemisphere, right column shows lateral and medial view of right hemisphere

Figure 8 shows the association of CNR and CT for two exemplary regions with contrary scanner effects (superior frontal and lateral occipital).

**Figure 8:**
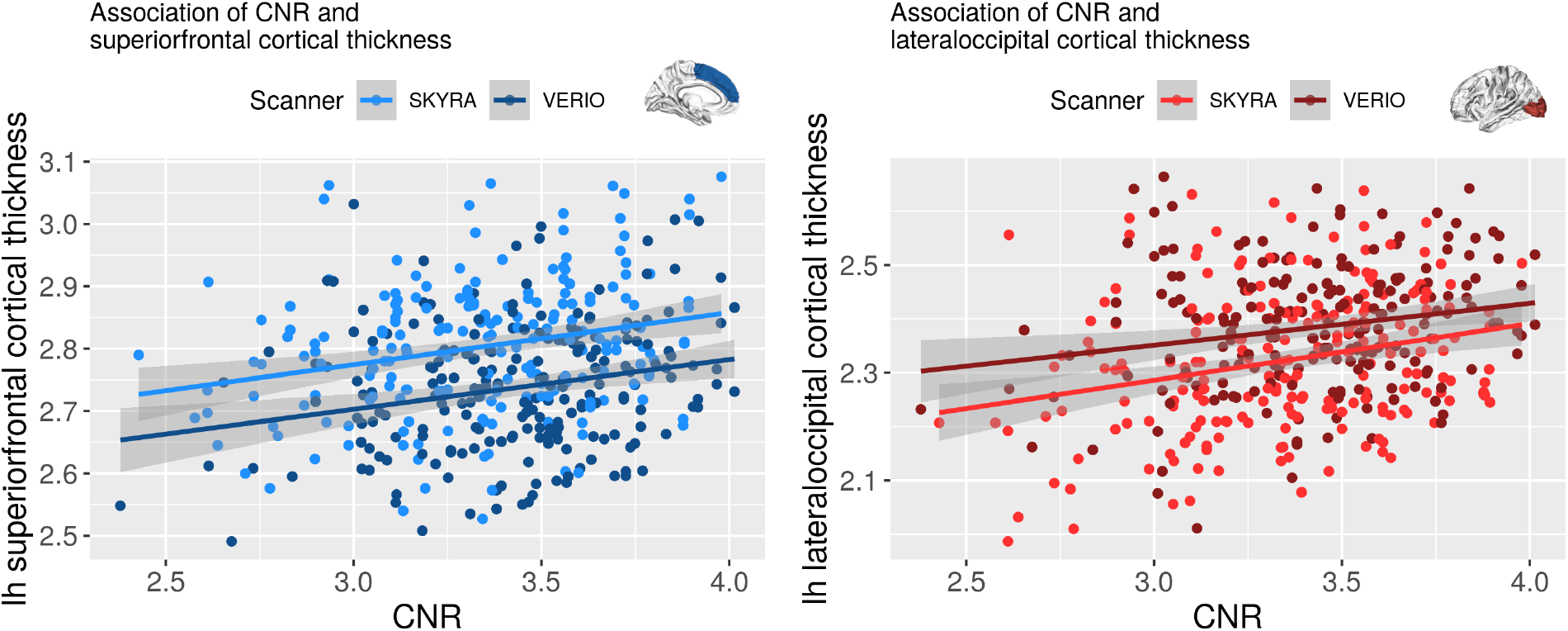
Association of CNR and cortical thickness in left superior frontal (left panel) and lateral occipital cortex (right panel)

### 4.4 Effect of offline gradient distortion correction

We examined whether the differences in cortical and subcortical GM measures arise from the difference in gradient distortion between the two scanners. We corrected both ND files using vendor-provided information on gradient distortions using `gradunwarp`.

Figure 9 shows the results for CT derived from the `gradunwarp` distortion corrected data (also see Table 5.1 in the Supplementary Material). The ICC was excellent throughout all ROI (mean= 0.91, min=0.8, max=0.98), and as expected, it was higher for the gradient distortion corrected data compared to the previous analysis of Verio ND vs Skyra D (mean ICC `gradunwarp` Skyra D vs Verio D: 0.91, mean ICC Skyra D vs Verio ND: 0.89, paired t-test: T = −4.04, p < 0.001).

**Figure 9:**
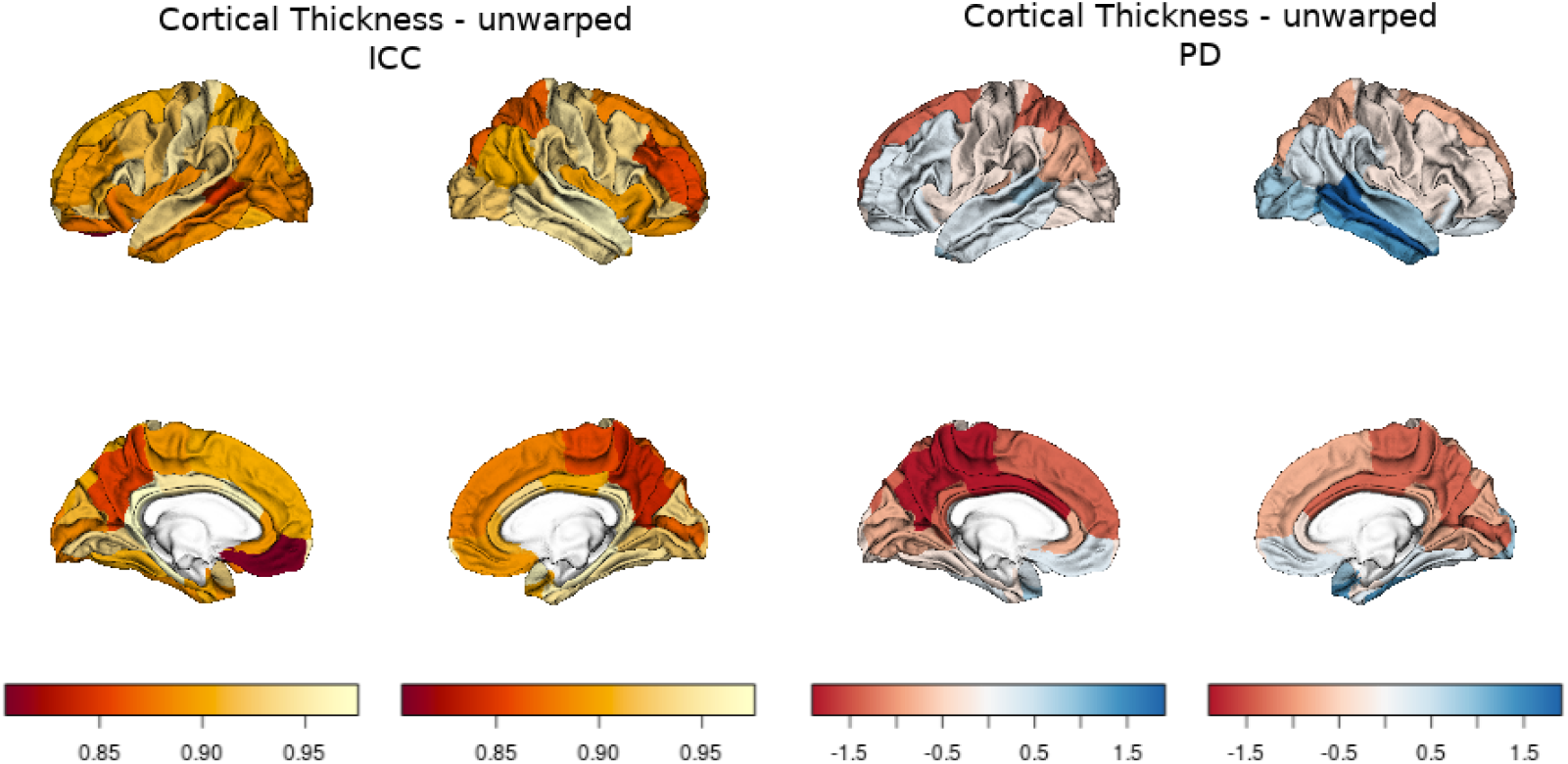
Comparison of CT results from gradunwarp-corrected data. Left panel: CT ICC, right panel: CT PD (for each panel, left column shows lateral and medial view of left hemisphere, right column shows lateral and medial view of right hemisphere), negative values: Skyra>Verio, positive values: Verio>Skyra

PD was around 1-3% (mean=-0.25%, min=-1.79%, max=1.89), with the same frontal-occipital pattern of biases. Accordingly, a paired t-test shows that the systematic differences between scanners slightly decreased after gradient distortion correction (mean PD `gradunwarp` Skyra D vs Verio D: 0.63, mean PD Skyra D vs Verio ND: 0.92, paired t-test: T = 3.92, p < 0.001). Yet, there were still significant differences after `gradunwarp` for most regions of interest (FDR-corrected, 62.5% of 64 bilateral cortical ROIs).

Table 2 shows the results for subcortical volumes derived from the gradient distortion corrected data. The ICC is excellent in all regions, similar to the cortical analysis (mean=0.95, min=0.81, max=0.99). For subcortical volumes, gradient distortion correction did not lead to a further improvement in ICC (mean ICC `gradunwarp` Skyra D vs Verio D = 0.95, mean ICC Skyra D vs Verio ND = 0.95, paired t-test: T = 0.81, p= 0.43).

**Table 2:** Reliability and percent difference for subcortical volumes from gradient non-linearity corrected data (T<0 reflects Skyra>Verio, T>0 reflects Verio>Skyra)

| ROI | hemi | ICC | lower ICC | upper ICC | PD | T | p | adj.p |
| --- | --- | --- | --- | --- | --- | --- | --- | --- |
| Thalamus | Left | 0.97 | 0.97 | 0.97 | 1.93 | -13.33 | 0.00 | **0** |
| Thalamus | Right | 0.98 | 0.97 | 0.98 | 1.64 | -10.98 | 0.00 | **0** |
| Caudate | Left | 0.99 | 0.99 | 0.99 | 1.46 | -8.34 | 0.00 | **0** |
| Caudate | Right | 0.99 | 0.99 | 0.99 | 1.78 | -13.43 | 0.00 | **0** |
| Putamen | Left | 0.98 | 0.97 | 0.98 | 1.63 | -7.12 | 0.00 | **0** |
| Putamen | Right | 0.99 | 0.98 | 0.99 | 1.39 | -8.57 | 0.00 | **0** |
| Pallidum | Left | 0.97 | 0.96 | 0.98 | 2.49 | -2.92 | 0.00 | **0** |
| Pallidum | Right | 0.94 | 0.92 | 0.96 | 2.87 | -2.29 | 0.02 | **0.02** |
| Hippocampus | Left | 0.95 | 0.94 | 0.96 | 2.29 | -13.95 | 0.00 | **0** |
| Hippocampus | Right | 0.96 | 0.95 | 0.97 | 1.91 | -10.35 | 0.00 | **0** |
| Amygdala | Left | 0.91 | 0.88 | 0.93 | 3.58 | -2.78 | 0.01 | **0.01** |
| Amygdala | Right | 0.94 | 0.91 | 0.95 | 2.99 | -2.93 | 0.00 | **0** |
| Accumbens | Left | 0.81 | 0.78 | 0.85 | 9.13 | -8.72 | 0.00 | **0** |
| Accumbens | Right | 0.94 | 0.93 | 0.95 | 4.11 | -3.94 | 0.00 | **0** |

The PD was around 2-3% (mean=2.8%, min=1.39%, max=9.13%) and did not differ from the original analysis (mean PD `gradunwarp` Skyra D vs Verio D = 2.8%, mean PD Skyra D vs Verio ND = 2.77%, paired t-test: T= −0.59, p= 0.57). There were significant differences after `gradunwarp` for all regions of interest (FDR-corrected, 100% of 14 bilateral subcortical regions).

## 5 Discussion

### Summary

In this paper, we aimed to investigate the reliability and bias in GM structure induced by a scanner upgrade in a longitudinal study. We compared outcomes of FreeSurfer’s longitudinal pipeline between two different MRI scanners with subsequent versions. We found between-scanner reliability measured with ICC to be excellent. Yet, paired t-tests revealed statistically significant differences, i.e. biases, in GM volume, area and thickness for a large number of cortical and subcortical regions. Offline correction for gradient distortions based on vendor-provided gradient information only slightly reduced these differences. T1-imaging based quality measures differed systematically between scanners, also when adjusting for gradient distortions. We conclude that scanner upgrades during a longitudinal study introduce bias in measures of cortical and subcortical grey matter structure and make it difficult to detect true effects when these are subtle like in the case of healthy aging, e.g. ~ 1% annual hippocampal volume loss in older healthy adults (Fraser, Shaw, and Cherbuin 2015). Therefore, before upgrading a MRI system during an ongoing longitudinal study, researchers should prepare to implement an appropriate correction method, such as deriving scaling factors from repeated measures before/after the upgrade or statistical adjustment methods.

### Comparison to previous upgrade studies

The results of our study are in line with previous findings which have indicated systematic effects of scanner upgrade on GM imaging outcomes (Lee et al. 2019; Han et al. 2006; Jovicich et al. 2009; Brunton et al. 2015).

Notably, in two recent studies, an upgrade from Magnetom Trio to Prismafit induced a significant increase in cortical thickness (CT) and volume as well as differences in subcortical volumes (Potvin et al. 2019; Plitman et al. 2020). Similar to our findings, ICC values for cortical measures were good to excellent in both studies. While the size of biases was comparable to our results (around 1-5% for cortical PD for CV and CT in (Potvin et al. 2019)), the location of the biased regions was different. We found a pattern of frontal to occipital differences, with frontal-precentral regions biased towards higher CT and GM volume values in Skyra compared to Verio, and occipital regions biased towards higher CT and GM volume in Verio. Yet, in (Potvin et al. 2019; Plitman et al. 2020) the biased regions were located in prefrontal and temporal regions where CT and GM volume consistently increased with the upgrade.

As shown above, there is a mismatch in location and direction of the upgrade effect between our study and (Potvin et al. 2019; Plitman et al. 2020). Thus, additional factors in our studies probably led to the observed frontal-occipital bias, and contributed to higher CT and GM volumes in Skyra compared to Verio.

One contributing factor is the observed image quality differences between the scanners. CNR and other measures of image quality indicated higher quality on the (earlier) Verio scanner. In contrast, studies investigating the effects of real upgrades showed increased image quality (quantified as signal-to-noise ratio (SNR) or CNR) after the upgrade (Potvin et al. 2019; Plitman et al. 2020). This may contribute to the observed bias pattern in GMV, yet increased SNR/CNR was also associated with higher CT in our study, independent of scanner-dependent regional CT differences (Shuter et al. 2008; Potvin et al. 2019).

Another possible factor contributing to the bias might be gradient distortion correction, which was different between acquisitions in the main analysis. According to (Jovicich et al. 2006), gradient distortions may account for 16% of the image intensity relative error, and adjusting for these distortions has previously removed site-related variations to <1% and increased reliability of the between-site scans to within-site level (Cannon et al. 2014). Yet, we did not see a substantial reduction of bias when applying offline gradient distortion correction. While for cortical measures, ICC slightly increased and PD decreased, subcortical volumes measures did not change. This is expected, as gradient distortions have most pronounced effects at the edges of the image; therefore their correction will affect objects at the center less than objects in the periphery, leaving subcortical areas nearly unchanged.

Finally, scaling differences might have contributed to the bias pattern between scanners. This is supported by the frontal-to-occipital bias pattern, as well as visual inspections of the longitudinal runs (i.e. when both had been registered to a common template), where subtle expansion of the brain in Skyra compared to Verio was observed. Taken together, we believe the systematic biases between Verio and Skyra stem from both scaling and image quality differences, and were strongly related to scanner hardware.

While our results certainly overestimate the effects of a real upgrade as discussed above, they still support previous studies on the biasing effects of a scanner upgrade and urge for the use of an adequate correction method if an upgrade becomes necessary during a longitudinal study. One possibility is to measure the same subjects shortly before and after the upgrade and to derive scaling factors like in (Keshavan et al. 2016). Another possibility, which does not require additional data acquisition, is longitudinal ComBat correction, which takes into account biased mean and scaling due to systematic scanner differences (Beer et al. 2020).

### Limitations

The main limitation of our study is that we did not assess the impact of a true upgrade (i.e. repeated measurements on the same scanner), instead we performed a site-comparison in which the MRI scanners at the two sides were as similar as possible. Another limitation is that we did not acquire multiple scans on the same system, nor randomized the order of participants across scanners.

### Strengths

Our study is well-powered, which is important to adequately compute reliability with an acceptable confidence interval. Also, we applied region-and brain-wide analyses, adjusted for gradient distortions and calculated complementary measures of reliability. Additionally, we present quantitative quality control measures derived from mriqc, a state-of-the-art quality control software.

### Conclusions

Taken together, in this study, we investigated the impact of a scanner upgrade on longitudinal cortical and subcortical GM measures. We found high reliability but strong regional biases in most regions of interest. While we possibly overestimated the effects of a real upgrade, this study urges for careful monitoring of scanner upgrades and adjustment of biases in longitudinal imaging studies. This may be achieved by deriving scaling factors immediately before/after the upgrade or by using longitudinal batch correction.

