## Supplemental Material for "Estimating the effect of a scanner upgrade on measures of grey matter structure for longitudinal designs"

**4 Correlation of CNR and CT** **7**

**5 Cortical thickness reliability and percent difference table for gradunwarp distortion corrected data** **9**

### 1 Cortical thickness

#### 1.1 Vertex-wise analysis of cortical thickness ICC

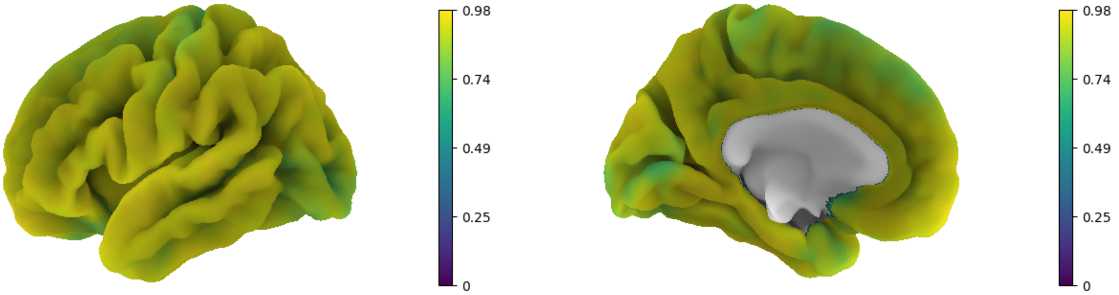

Figure 1: Lateral and medial view of ICC for thickness (left, right panel)

#### 1.2 Vertex-wise analysis of cortical thickness PD

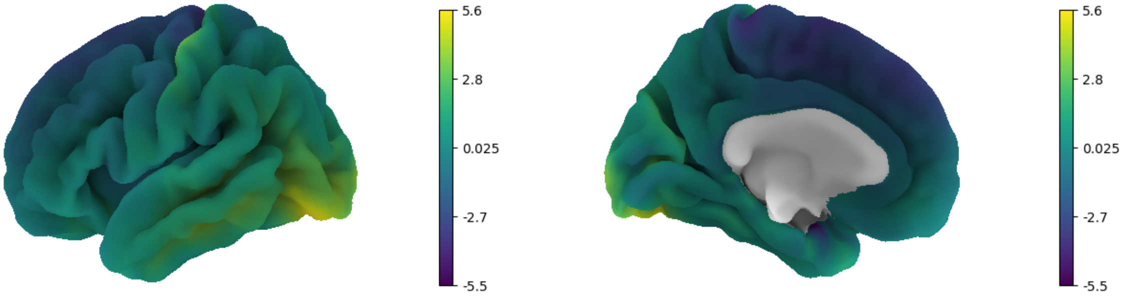

Figure 2: Lateral and medial view of PD for thickness (positive values: Verio>Skyra , negative values: Verio<Skyra ))

#### 1.3 Reliability and percent difference table of cortical thickness

Table 1: Reliability and differences for cortical thickness

| ROI | hemi | ICC | lower ICC | upper ICC | PD | T | p | adj.p |
|---|---|---|---|---|---|---|---|---|
| bankssts | lh | 0.841 | 0.772 | 0.850 | 1.97 | 9.88 | 0.00 | **0** |
| bankssts | rh | 0.909 | 0.904 | 0.936 | 1.47 | 8.92 | 0.00 | **0** |
| caudalanteriorcingulate | lh | 0.958 | 0.956 | 0.967 | -1.20 | -6.60 | 0.00 | **0** |
| caudalanteriorcingulate | rh | 0.880 | 0.882 | 0.901 | -2.45 | -11.47 | 0.00 | **0** |
| caudalmiddlefrontal | lh | 0.884 | 0.859 | 0.904 | -0.91 | -5.31 | 0.00 | **0** |
| caudalmiddlefrontal | rh | 0.895 | 0.859 | 0.913 | -0.37 | -2.16 | 0.03 | **0.04** |
| cuneus | lh | 0.884 | 0.831 | 0.898 | 1.25 | 4.85 | 0.00 | **0** |
| cuneus | rh | 0.919 | 0.895 | 0.925 | 0.42 | 1.70 | 0.09 | 0.12 |
| entorhinal | lh | 0.916 | 0.870 | 0.929 | -0.15 | -0.51 | 0.61 | 0.67 |
| entorhinal | rh | 0.937 | 0.919 | 0.940 | -0.19 | -0.69 | 0.49 | 0.55 |
| fusiform | lh | 0.883 | 0.863 | 0.901 | 0.96 | 6.00 | 0.00 | **0** |
| fusiform | rh | 0.929 | 0.923 | 0.940 | 0.42 | 3.37 | 0.00 | **0** |
| inferiorparietal | lh | 0.863 | 0.831 | 0.890 | 1.21 | 7.78 | 0.00 | **0** |
| inferiorparietal | rh | 0.930 | 0.904 | 0.930 | 0.39 | 2.88 | 0.00 | **0** |
| inferiortemporal | lh | 0.858 | 0.857 | 0.908 | 1.74 | 10.85 | 0.00 | **0** |
| inferiortemporal | rh | 0.910 | 0.910 | 0.925 | 1.07 | 8.40 | 0.00 | **0** |
| isthmuscingulate | lh | 0.979 | 0.973 | 0.984 | -0.39 | -2.50 | 0.01 | **0.01** |
| isthmuscingulate | rh | 0.950 | 0.929 | 0.953 | -0.96 | -5.47 | 0.00 | **0** |
| lateraloccipital | lh | 0.742 | 0.695 | 0.780 | 3.23 | 15.52 | 0.00 | **0** |
| lateraloccipital | rh | 0.857 | 0.822 | 0.855 | 1.86 | 9.34 | 0.00 | **0** |
| lateralorbitofrontal | lh | 0.860 | 0.832 | 0.887 | 0.73 | 4.24 | 0.00 | **0** |
| lateralorbitofrontal | rh | 0.794 | 0.768 | 0.823 | 0.65 | 2.42 | 0.02 | **0.03** |
| lingual | lh | 0.918 | 0.904 | 0.940 | 0.78 | 3.99 | 0.00 | **0** |
| lingual | rh | 0.923 | 0.901 | 0.943 | 0.61 | 3.21 | 0.00 | **0** |
| medialorbitofrontal | lh | 0.849 | 0.846 | 0.871 | 1.07 | 4.12 | 0.00 | **0** |
| medialorbitofrontal | rh | 0.870 | 0.847 | 0.926 | -0.08 | -0.37 | 0.71 | 0.74 |
| middletemporal | lh | 0.875 | 0.850 | 0.899 | 1.76 | 11.94 | 0.00 | **0** |
| middletemporal | rh | 0.905 | 0.899 | 0.930 | 1.26 | 9.41 | 0.00 | **0** |
| parahippocampal | lh | 0.964 | 0.954 | 0.972 | 0.73 | 3.87 | 0.00 | **0** |
| parahippocampal | rh | 0.962 | 0.959 | 0.970 | 0.02 | -0.02 | 0.99 | 1 |
| paracentral | lh | 0.824 | 0.806 | 0.843 | -2.34 | -12.51 | 0.00 | **0** |
| paracentral | rh | 0.710 | 0.605 | 0.712 | -3.16 | -13.62 | 0.00 | **0** |
| parsopercularis | lh | 0.949 | 0.938 | 0.957 | -0.16 | -1.30 | 0.20 | 0.25 |
| parsopercularis | rh | 0.935 | 0.904 | 0.949 | 0.07 | 0.45 | 0.65 | 0.7 |
| parsorbitalis | lh | 0.930 | 0.914 | 0.944 | 0.29 | 1.62 | 0.11 | 0.14 |
| parsorbitalis | rh | 0.942 | 0.918 | 0.944 | 0.20 | 0.95 | 0.34 | 0.41 |
| parstriangularis | lh | 0.913 | 0.880 | 0.945 | 0.28 | 1.90 | 0.06 | 0.08 |
| parstriangularis | rh | 0.901 | 0.885 | 0.932 | 0.36 | 2.02 | 0.05 | 0.07 |
| pericalcarine | lh | 0.838 | 0.835 | 0.908 | 1.65 | 4.77 | 0.00 | **0** |
| pericalcarine | rh | 0.796 | 0.718 | 0.803 | 2.42 | 5.93 | 0.00 | **0** |
| postcentral | lh | 0.925 | 0.904 | 0.949 | 0.94 | 5.78 | 0.00 | **0** |
| postcentral | rh | 0.929 | 0.904 | 0.948 | 0.98 | 6.50 | 0.00 | **0** |
| posteriorcingulate | lh | 0.924 | 0.886 | 0.929 | -1.55 | -9.58 | 0.00 | **0** |
| posteriorcingulate | rh | 0.819 | 0.756 | 0.832 | -2.43 | -13.68 | 0.00 | **0** |
| precentral | lh | 0.893 | 0.873 | 0.909 | -1.03 | -7.25 | 0.00 | **0** |
| precentral | rh | 0.933 | 0.907 | 0.945 | -0.73 | -4.85 | 0.00 | **0** |
| precuneus | lh | 0.910 | 0.870 | 0.910 | 0.00 | -0.04 | 0.97 | 1 |
| precuneus | rh | 0.884 | 0.884 | 0.895 | -0.86 | -5.63 | 0.00 | **0** |
| rostralanteriorcingulate | lh | 0.814 | 0.776 | 0.886 | -0.42 | -1.36 | 0.18 | 0.23 |
| rostralanteriorcingulate | rh | 0.920 | 0.906 | 0.937 | -0.80 | -4.24 | 0.00 | **0** |
| rostralmiddlefrontal | lh | 0.892 | 0.870 | 0.924 | -0.53 | -3.41 | 0.00 | **0** |
| rostralmiddlefrontal | rh | 0.857 | 0.837 | 0.886 | 0.00 | 0.00 | 1.00 | 1 |
| superiorfrontal | lh | 0.805 | 0.749 | 0.790 | -2.01 | -13.08 | 0.00 | **0** |
| superiorfrontal | rh | 0.687 | 0.671 | 0.739 | -2.67 | -16.08 | 0.00 | **0** |
| superiorparietal | lh | 0.925 | 0.909 | 0.927 | 0.43 | 2.84 | 0.01 | **0.01** |

### 2 Cortical area

#### 2.1 Vertex-wise analysis of cortical area ICC

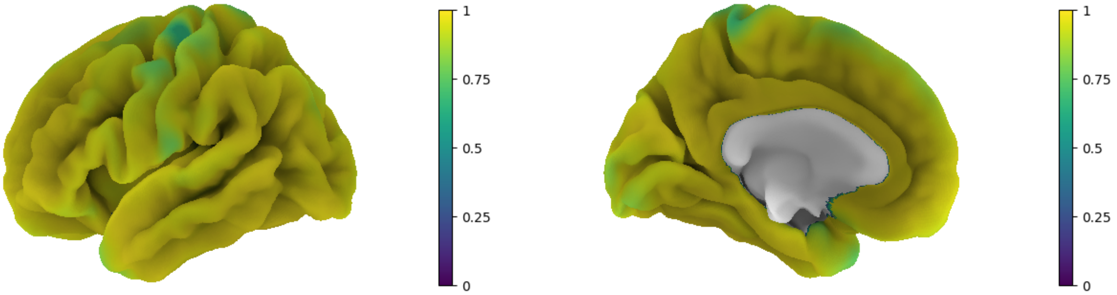

Figure 3: Lateral and medial view of ICC for area (left, right panel)

#### 2.2 Vertex-wise analysis of cortical area PD

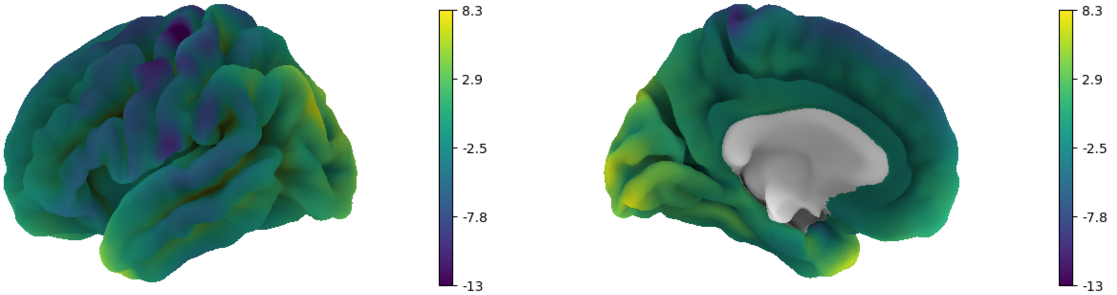

Figure 4: Lateral and medial view of PD for area (positive values: Verio>Skyra , negative values: Verio<Skyra )

#### 2.3 Reliability and percent difference table of cortical area

Table 2: Reliability and differences for cortical area

| ROI | hemi | ICC | lower ICC | upper ICC | PD | T | p | adj.p |
|---|---|---|---|---|---|---|---|---|
| bankssts | lh | 0.988 | 0.986 | 0.988 | 2.39 | 21.51 | 0.00 | **0** |
| bankssts | rh | 0.981 | 0.971 | 0.983 | 2.69 | 23.79 | 0.00 | **0** |
| caudalanteriorcingulate | lh | 0.995 | 0.995 | 0.995 | -1.07 | -6.56 | 0.00 | **0** |
| caudalanteriorcingulate | rh | 0.992 | 0.984 | 0.993 | -1.41 | -6.94 | 0.00 | **0** |
| caudalmiddlefrontal | lh | 0.993 | 0.992 | 0.994 | -1.26 | -11.35 | 0.00 | **0** |
| caudalmiddlefrontal | rh | 0.994 | 0.992 | 0.995 | -1.12 | -8.90 | 0.00 | **0** |
| cuneus | lh | 0.991 | 0.990 | 0.993 | 1.22 | 7.61 | 0.00 | **0** |
| cuneus | rh | 0.991 | 0.988 | 0.991 | 0.92 | 5.39 | 0.00 | **0** |
| entorhinal | lh | 0.981 | 0.977 | 0.984 | -0.73 | -2.03 | 0.04 | 0.05 |
| entorhinal | rh | 0.977 | 0.955 | 0.982 | 0.23 | 0.65 | 0.52 | 0.54 |
| fusiform | lh | 0.996 | 0.996 | 0.997 | 0.30 | 3.19 | 0.00 | **0** |
| fusiform | rh | 0.996 | 0.994 | 0.997 | 0.48 | 5.36 | 0.00 | **0** |
| inferiorparietal | lh | 0.984 | 0.980 | 0.988 | 1.94 | 12.81 | 0.00 | **0** |
| inferiorparietal | rh | 0.988 | 0.985 | 0.990 | 1.54 | 10.76 | 0.00 | **0** |
| inferiortemporal | lh | 0.997 | 0.997 | 0.998 | 0.28 | 3.19 | 0.00 | **0** |
| inferiortemporal | rh | 0.997 | 0.996 | 0.997 | 0.55 | 5.67 | 0.00 | **0** |
| isthmuscingulate | lh | 0.994 | 0.992 | 0.995 | 0.13 | 0.85 | 0.40 | 0.43 |
| isthmuscingulate | rh | 0.988 | 0.985 | 0.990 | -0.49 | -2.40 | 0.02 | **0.02** |
| lateraloccipital | lh | 0.975 | 0.976 | 0.979 | 2.30 | 14.74 | 0.00 | **0** |
| lateraloccipital | rh | 0.976 | 0.970 | 0.976 | 2.23 | 13.75 | 0.00 | **0** |
| lateralorbitofrontal | lh | 0.993 | 0.991 | 0.997 | 0.32 | 2.69 | 0.01 | **0.01** |
| lateralorbitofrontal | rh | 0.944 | 0.921 | 0.952 | -0.88 | -2.63 | 0.01 | **0.01** |
| lingual | lh | 0.980 | 0.977 | 0.983 | 2.08 | 13.31 | 0.00 | **0** |
| lingual | rh | 0.988 | 0.983 | 0.990 | 1.25 | 7.89 | 0.00 | **0** |
| medialorbitofrontal | lh | 0.926 | 0.888 | 0.938 | -1.55 | -3.44 | 0.00 | **0** |
| medialorbitofrontal | rh | 0.944 | 0.921 | 0.946 | -1.41 | -4.63 | 0.00 | **0** |
| middletemporal | lh | 0.996 | 0.994 | 0.997 | -0.48 | -3.92 | 0.00 | **0** |
| middletemporal | rh | 0.997 | 0.996 | 0.998 | -0.18 | -1.93 | 0.06 | 0.07 |
| parahippocampal | lh | 0.986 | 0.981 | 0.988 | -0.04 | -0.32 | 0.75 | 0.76 |
| parahippocampal | rh | 0.990 | 0.986 | 0.992 | -0.66 | -4.83 | 0.00 | **0** |
| paracentral | lh | 0.980 | 0.972 | 0.984 | -1.75 | -11.07 | 0.00 | **0** |
| paracentral | rh | 0.976 | 0.964 | 0.980 | -1.86 | -10.97 | 0.00 | **0** |
| parsopercularis | lh | 0.996 | 0.995 | 0.996 | -0.66 | -6.33 | 0.00 | **0** |
| parsopercularis | rh | 0.995 | 0.991 | 0.995 | -0.28 | -2.49 | 0.01 | **0.01** |
| parsorbitalis | lh | 0.969 | 0.962 | 0.973 | -2.57 | -15.40 | 0.00 | **0** |
| parsorbitalis | rh | 0.982 | 0.977 | 0.986 | -1.69 | -10.23 | 0.00 | **0** |
| parstriangularis | lh | 0.994 | 0.993 | 0.995 | -1.10 | -9.89 | 0.00 | **0** |
| parstriangularis | rh | 0.994 | 0.992 | 0.994 | -1.38 | -11.42 | 0.00 | **0** |
| pericalcarine | lh | 0.994 | 0.992 | 0.994 | 1.07 | 6.90 | 0.00 | **0** |
| pericalcarine | rh | 0.994 | 0.993 | 0.995 | 0.40 | 2.36 | 0.02 | **0.02** |
| postcentral | lh | 0.967 | 0.968 | 0.973 | -2.37 | -15.21 | 0.00 | **0** |
| postcentral | rh | 0.963 | 0.953 | 0.968 | -2.58 | -15.13 | 0.00 | **0** |
| posteriorcingulate | lh | 0.995 | 0.994 | 0.996 | -0.57 | -4.58 | 0.00 | **0** |
| posteriorcingulate | rh | 0.992 | 0.988 | 0.993 | -0.79 | -6.08 | 0.00 | **0** |
| precentral | lh | 0.973 | 0.967 | 0.977 | -2.09 | -15.52 | 0.00 | **0** |
| precentral | rh | 0.977 | 0.971 | 0.978 | -1.76 | -13.39 | 0.00 | **0** |
| precuneus | lh | 0.995 | 0.993 | 0.996 | -0.36 | -3.17 | 0.00 | **0** |
| precuneus | rh | 0.995 | 0.994 | 0.996 | -0.42 | -3.74 | 0.00 | **0** |
| rostralanteriorcingulate | lh | 0.972 | 0.958 | 0.983 | -1.45 | -3.93 | 0.00 | **0** |
| rostralanteriorcingulate | rh | 0.981 | 0.976 | 0.986 | -2.20 | -7.33 | 0.00 | **0** |
| rostralmiddlefrontal | lh | 0.996 | 0.995 | 0.996 | -0.69 | -7.16 | 0.00 | **0** |
| rostralmiddlefrontal | rh | 0.994 | 0.989 | 0.996 | -0.60 | -4.44 | 0.00 | **0** |
| superiorfrontal | lh | 0.975 | 0.970 | 0.980 | -2.22 | -19.09 | 0.00 | **0** |
| superiorfrontal | rh | 0.981 | 0.979 | 0.985 | -2.07 | -18.02 | 0.00 | **0** |
| superiorparietal | lh | 0.992 | 0.991 | 0.993 | 0.54 | 3.55 | 0.00 | **0** |

### 3 Cortical volume

#### 3.1 Vertex-wise analysis of cortical volume ICC

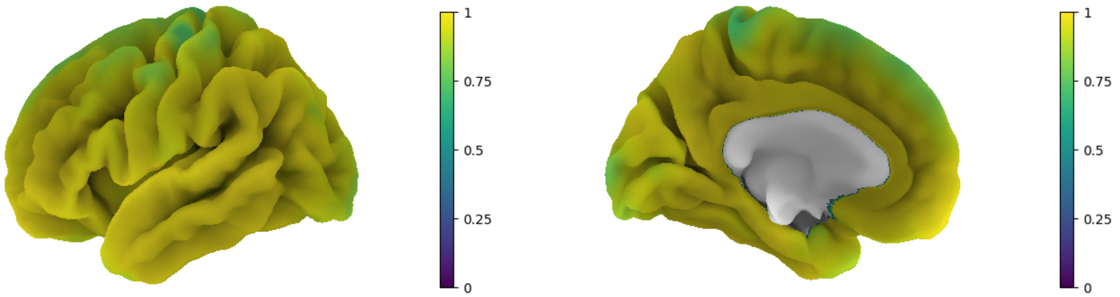

Figure 5: Lateral and medial view of ICC for volume (left, right panel)

#### 3.2 Vertex-wise analysis of cortical volume PD

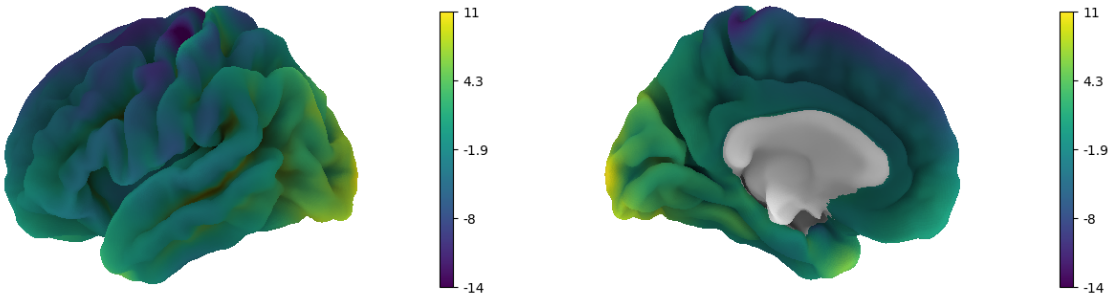

Figure 6: Lateral and medial view of PD for volume (positive values: Verio>Skyra , negative values: Verio<Skyra )

#### 3.3 Reliability and percent difference table of cortical volume

Table 3: Reliability and differences for cortical volume

| ROI | hemi | ICC | lower ICC | upper ICC | PD | T | p | adj.p |
|---|---|---|---|---|---|---|---|---|
| Thalamus | Left | 0.97 | 0.97 | 0.98 | 1.90 | -12.27 | 0.00 | **0** |
| Thalamus | Right | 0.98 | 0.97 | 0.99 | 1.61 | -9.37 | 0.00 | **0** |
| Caudate | Left | 0.99 | 0.99 | 0.99 | 1.65 | -10.64 | 0.00 | **0** |
| Caudate | Right | 0.98 | 0.98 | 0.99 | 2.12 | -15.72 | 0.00 | **0** |
| Putamen | Left | 0.98 | 0.98 | 0.99 | 1.58 | -6.22 | 0.00 | **0** |
| Putamen | Right | 0.99 | 0.99 | 0.99 | 1.22 | -4.85 | 0.00 | **0** |
| Pallidum | Left | 0.96 | 0.95 | 0.97 | 2.48 | -0.44 | 0.66 | 0.66 |
| Pallidum | Right | 0.95 | 0.95 | 0.96 | 2.53 | -3.58 | 0.00 | **0** |
| Hippocampus | Left | 0.96 | 0.95 | 0.97 | 2.01 | -8.51 | 0.00 | **0** |
| Hippocampus | Right | 0.97 | 0.97 | 0.97 | 1.61 | -8.16 | 0.00 | **0** |
| Amygdala | Left | 0.91 | 0.89 | 0.92 | 3.57 | -1.47 | 0.14 | 0.15 |
| Amygdala | Right | 0.94 | 0.92 | 0.96 | 3.00 | -2.73 | 0.01 | **0.01** |
| Accumbens | Left | 0.81 | 0.74 | 0.85 | 9.52 | -9.30 | 0.00 | **0** |
| Accumbens | Right | 0.95 | 0.93 | 0.96 | 3.91 | -5.53 | 0.00 | **0** |

### 4 Correlation of CNR and CT

Table 4: Association of CNR and scanner with CT

| ROI | hemi | CNR estimate | scanner estimate | p CNR | p scanner | adj.p.CNR | adj.p.scanner |
|---|---|---|---|---|---|---|---|
| bankssts | lh | 0.068 | 0.044 | 0.000 | 0.00 | **0.001** | **0** |
| bankssts | rh | 0.064 | 0.034 | 0.000 | 0.00 | **0.005** | **0.01** |
| caudalanteriorcingulate | lh | 0.035 | -0.036 | 0.000 | 0.00 | 0.133 | 0.16 |
| caudalanteriorcingulate | rh | -0.028 | -0.060 | 0.000 | 0.00 | 0.31 | 0.33 |
| caudalmiddlefrontal | lh | 0.050 | -0.029 | 0.000 | 0.00 | **0.007** | **0.02** |
| caudalmiddlefrontal | rh | 0.059 | -0.016 | 0.006 | 0.01 | **0.008** | **0.02** |
| cuneus | lh | 0.058 | 0.020 | 0.000 | 0.00 | **0.005** | **0.01** |
| cuneus | rh | 0.046 | 0.011 | 0.067 | 0.10 | 0.06 | 0.09 |
| entorhinal | lh | 0.073 | -0.011 | 0.272 | 0.34 | 0.071 | 0.09 |
| entorhinal | rh | 0.071 | -0.010 | 0.404 | 0.47 | 0.162 | 0.19 |
| fusiform | lh | 0.078 | 0.020 | 0.000 | 0.00 | **0** | **0** |
| fusiform | rh | 0.044 | 0.008 | 0.088 | 0.12 | **0.016** | **0.03** |
| inferiorparietal | lh | 0.070 | 0.024 | 0.000 | 0.00 | **0** | **0** |
| inferiorparietal | rh | 0.031 | 0.008 | 0.066 | 0.10 | 0.092 | 0.12 |
| inferiortemporal | lh | 0.065 | 0.043 | 0.000 | 0.00 | **0.001** | **0** |
| inferiortemporal | rh | 0.029 | 0.028 | 0.000 | 0.00 | 0.14 | 0.16 |
| isthmuscingulate | lh | 0.028 | -0.011 | 0.004 | 0.01 | 0.095 | 0.12 |
| isthmuscingulate | rh | 0.051 | -0.029 | 0.000 | 0.00 | **0.03** | 0.05 |
| lateraloccipital | lh | 0.078 | 0.067 | 0.000 | 0.00 | **0** | **0** |
| lateraloccipital | rh | 0.054 | 0.043 | 0.000 | 0.00 | **0.022** | **0.04** |
| lateralorbitofrontal | lh | 0.052 | 0.015 | 0.003 | 0.00 | **0.004** | **0.01** |
| lateralorbitofrontal | rh | 0.052 | 0.011 | 0.182 | 0.23 | 0.063 | 0.09 |
| lingual | lh | 0.070 | 0.011 | 0.016 | 0.03 | **0** | **0** |
| lingual | rh | 0.044 | 0.014 | 0.008 | 0.01 | **0.036** | 0.06 |
| medialorbitofrontal | lh | -0.002 | 0.026 | 0.000 | 0.00 | 0.918 | 0.92 |
| medialorbitofrontal | rh | 0.030 | -0.004 | 0.545 | 0.59 | 0.239 | 0.27 |
| middletemporal | lh | 0.056 | 0.047 | 0.000 | 0.00 | **0.003** | **0.01** |
| middletemporal | rh | 0.052 | 0.033 | 0.000 | 0.00 | **0.015** | **0.03** |
| parahippocampal | lh | 0.069 | 0.015 | 0.011 | 0.02 | **0.005** | **0.01** |
| parahippocampal | rh | 0.055 | -0.007 | 0.314 | 0.38 | 0.052 | 0.07 |
| paracentral | lh | 0.049 | -0.064 | 0.000 | 0.00 | **0.013** | **0.03** |
| paracentral | rh | 0.012 | -0.087 | 0.000 | 0.00 | 0.663 | 0.68 |
| parsopercularis | lh | 0.043 | -0.008 | 0.023 | 0.04 | **0.003** | **0.01** |
| parsopercularis | rh | 0.023 | -0.002 | 0.746 | 0.77 | 0.243 | 0.27 |
| parsorbitalis | lh | 0.060 | 0.003 | 0.591 | 0.63 | **0.005** | **0.01** |
| parsorbitalis | rh | 0.062 | -0.002 | 0.785 | 0.80 | **0.015** | **0.03** |
| parstriangularis | lh | 0.046 | 0.003 | 0.416 | 0.48 | **0.005** | **0.01** |
| parstriangularis | rh | 0.050 | 0.003 | 0.481 | 0.54 | **0.013** | **0.03** |
| pericalcarine | lh | 0.049 | 0.026 | 0.000 | 0.00 | **0.042** | 0.06 |
| pericalcarine | rh | 0.073 | 0.047 | 0.000 | 0.00 | **0.035** | 0.05 |
| postcentral | lh | 0.058 | 0.016 | 0.000 | 0.00 | **0** | **0** |
| postcentral | rh | 0.051 | 0.018 | 0.000 | 0.00 | **0.002** | **0.01** |
| posteriorcingulate | lh | 0.046 | -0.043 | 0.000 | 0.00 | **0.013** | **0.03** |
| posteriorcingulate | rh | -0.007 | -0.061 | 0.000 | 0.00 | 0.754 | 0.77 |
| precentral | lh | 0.060 | -0.034 | 0.000 | 0.00 | **0** | **0** |
| precentral | rh | 0.062 | -0.027 | 0.000 | 0.00 | **0.002** | **0.01** |
| precuneus | lh | 0.059 | -0.005 | 0.187 | 0.24 | **0** | **0** |
| precuneus | rh | 0.041 | -0.023 | 0.000 | 0.00 | **0.028** | 0.05 |
| rostralanteriorcingulate | lh | 0.117 | -0.023 | 0.023 | 0.04 | **0** | **0** |
| rostralanteriorcingulate | rh | -0.021 | -0.026 | 0.000 | 0.00 | 0.425 | 0.45 |
| rostralmiddlefrontal | lh | 0.048 | -0.017 | 0.000 | 0.00 | **0.002** | **0.01** |
| rostralmiddlefrontal | rh | 0.025 | -0.002 | 0.705 | 0.74 | 0.207 | 0.23 |
| superiorfrontal | lh | 0.048 | -0.060 | 0.000 | 0.00 | **0.007** | **0.02** |
| superiorfrontal | rh | 0.039 | -0.081 | 0.000 | 0.00 | 0.065 | 0.09 |
| superiorparietal | lh | 0.047 | 0.006 | 0.128 | 0.17 | **0.001** | **0.01** |

#### 5 Cortical thickness reliability and percent difference table for gradunwarp distortion corrected data

Table 5: Reliability and percent difference for cortical thickness from gradient non-linearity corrected data (T<0 reflects Skyra >Verio , T>0 reflects Verio >Skyra )

| ROI | hemi | ICC | lower ICC | upper ICC | PD | T | p | adj.p |
|---|---|---|---|---|---|---|---|---|
| bankssts | lh | 0.849 | 0.818 | 0.883 | 1.89 | 9.39 | 0.00 | **0** |
| bankssts | rh | 0.939 | 0.936 | 0.947 | 0.90 | 5.92 | 0.00 | **0** |
| caudalanteriorcingulate | lh | 0.966 | 0.955 | 0.973 | -1.07 | -6.40 | 0.00 | **0** |
| caudalanteriorcingulate | rh | 0.937 | 0.932 | 0.951 | -1.68 | -10.98 | 0.00 | **0** |
| caudalmiddlefrontal | lh | 0.915 | 0.916 | 0.932 | -0.39 | -2.53 | 0.01 | **0.02** |
| caudalmiddlefrontal | rh | 0.937 | 0.933 | 0.947 | 0.01 | 0.05 | 0.96 | 0.96 |
| cuneus | lh | 0.911 | 0.882 | 0.932 | -0.89 | -3.87 | 0.00 | **0** |
| cuneus | rh | 0.935 | 0.920 | 0.937 | -0.97 | -5.00 | 0.00 | **0** |
| entorhinal | lh | 0.900 | 0.889 | 0.917 | -0.21 | -0.55 | 0.58 | 0.65 |
| entorhinal | rh | 0.957 | 0.943 | 0.966 | -0.12 | -0.59 | 0.56 | 0.63 |
| fusiform | lh | 0.907 | 0.881 | 0.923 | 0.24 | 1.53 | 0.13 | 0.2 |
| fusiform | rh | 0.942 | 0.923 | 0.945 | -0.32 | -2.70 | 0.01 | **0.02** |
| inferiorparietal | lh | 0.887 | 0.865 | 0.900 | 0.28 | 1.77 | 0.08 | 0.13 |
| inferiorparietal | rh | 0.905 | 0.868 | 0.933 | -0.87 | -6.21 | 0.00 | **0** |
| inferiortemporal | lh | 0.889 | 0.875 | 0.903 | 1.46 | 10.36 | 0.00 | **0** |
| inferiortemporal | rh | 0.947 | 0.934 | 0.958 | 0.17 | 1.39 | 0.17 | 0.25 |
| isthmuscingulate | lh | 0.974 | 0.971 | 0.982 | -0.85 | -5.57 | 0.00 | **0** |
| isthmuscingulate | rh | 0.952 | 0.941 | 0.966 | -1.09 | -6.66 | 0.00 | **0** |
| lateraloccipital | lh | 0.883 | 0.874 | 0.920 | 0.99 | 4.90 | 0.00 | **0** |
| lateraloccipital | rh | 0.924 | 0.916 | 0.953 | -0.17 | -0.98 | 0.33 | 0.41 |
| lateralorbitofrontal | lh | 0.874 | 0.859 | 0.897 | 0.47 | 2.63 | 0.01 | **0.02** |
| lateralorbitofrontal | rh | 0.892 | 0.851 | 0.912 | 0.30 | 1.35 | 0.18 | 0.26 |
| lingual | lh | 0.937 | 0.917 | 0.944 | -0.87 | -5.27 | 0.00 | **0** |
| lingual | rh | 0.925 | 0.914 | 0.934 | -0.74 | -4.10 | 0.00 | **0** |
| medialorbitofrontal | lh | 0.805 | 0.782 | 0.863 | 0.30 | 0.87 | 0.39 | 0.46 |
| medialorbitofrontal | rh | 0.892 | 0.854 | 0.885 | 0.19 | 0.92 | 0.36 | 0.43 |
| middletemporal | lh | 0.877 | 0.860 | 0.913 | 1.73 | 12.07 | 0.00 | **0** |
| middletemporal | rh | 0.949 | 0.943 | 0.964 | 0.38 | 3.05 | 0.00 | **0** |
| parahippocampal | lh | 0.975 | 0.967 | 0.978 | 0.22 | 1.43 | 0.15 | 0.22 |
| parahippocampal | rh | 0.964 | 0.963 | 0.973 | -0.29 | -1.69 | 0.09 | 0.15 |
| paracentral | lh | 0.893 | 0.869 | 0.917 | -1.33 | -8.04 | 0.00 | **0** |
| paracentral | rh | 0.857 | 0.808 | 0.884 | -1.79 | -11.38 | 0.00 | **0** |
| parsopercularis | lh | 0.946 | 0.939 | 0.956 | -0.04 | -0.30 | 0.77 | 0.82 |
| parsopercularis | rh | 0.946 | 0.939 | 0.960 | 0.04 | 0.22 | 0.82 | 0.86 |
| parsorbitalis | lh | 0.926 | 0.876 | 0.931 | -0.14 | -0.77 | 0.44 | 0.51 |
| parsorbitalis | rh | 0.932 | 0.927 | 0.938 | 0.02 | 0.08 | 0.94 | 0.95 |
| parstriangularis | lh | 0.909 | 0.877 | 0.927 | -0.02 | -0.09 | 0.93 | 0.95 |
| parstriangularis | rh | 0.914 | 0.881 | 0.924 | 0.09 | 0.53 | 0.59 | 0.65 |
| pericalcarine | lh | 0.884 | 0.860 | 0.895 | -1.06 | -3.50 | 0.00 | **0** |
| pericalcarine | rh | 0.866 | 0.853 | 0.873 | -0.84 | -2.19 | 0.03 | 0.05 |
| postcentral | lh | 0.949 | 0.940 | 0.960 | -0.14 | -0.95 | 0.34 | 0.41 |
| postcentral | rh | 0.948 | 0.938 | 0.964 | -0.45 | -3.19 | 0.00 | **0** |
| posteriorcingulate | lh | 0.948 | 0.941 | 0.963 | -1.35 | -10.57 | 0.00 | **0** |
| posteriorcingulate | rh | 0.903 | 0.875 | 0.914 | -1.72 | -13.91 | 0.00 | **0** |
| precentral | lh | 0.919 | 0.892 | 0.937 | -0.45 | -3.20 | 0.00 | **0** |
| precentral | rh | 0.935 | 0.893 | 0.961 | -0.19 | -1.52 | 0.13 | 0.2 |
| precuneus | lh | 0.852 | 0.819 | 0.866 | -1.22 | -7.42 | 0.00 | **0** |
| precuneus | rh | 0.837 | 0.813 | 0.874 | -1.54 | -10.38 | 0.00 | **0** |
| rostralanteriorcingulate | lh | 0.892 | 0.869 | 0.907 | -0.27 | -1.11 | 0.27 | 0.35 |
| rostralanteriorcingulate | rh | 0.908 | 0.899 | 0.930 | -0.63 | -3.10 | 0.00 | **0** |
| rostralmiddlefrontal | lh | 0.893 | 0.865 | 0.920 | -0.38 | -2.42 | 0.02 | **0.03** |
| rostralmiddlefrontal | rh | 0.855 | 0.809 | 0.843 | 0.27 | 1.58 | 0.12 | 0.19 |
| superiorfrontal | lh | 0.905 | 0.884 | 0.921 | -0.87 | -6.73 | 0.00 | **0** |
| superiorfrontal | rh | 0.880 | 0.854 | 0.905 | -1.12 | -8.46 | 0.00 | **0** |
